# A single-cell transcriptomic map of the *Xenopus* mesonephros reveals conserved nephron patterning across vertebrate kidney forms

**DOI:** 10.1101/2025.01.13.632850

**Authors:** Mark E. Corkins, Adrian Romero, MaryAnne A. Achieng, Nils O. Lindström, Rachel K. Miller

## Abstract

The vertebrate kidney develops through three sequential forms: the pronephros, mesonephros, and metanephros. These three forms are composed of segmented nephrons that mediate fluid filtration and solute homeostasis. In aquatic egg-laying vertebrates such as *Xenopus laevis*, the mesonephros serves as the final functional renal organ, whereas in mammals it is a transient embryonic structure that precedes metanephric development. Despite its central developmental and evolutionary significance, the cellular and transcriptional organization of the mesonephric kidney remains poorly defined at single-cell resolution.

Here, we mapped the cellular and transcriptional landscape of the *Xenopus laevis* mesonephros from NF stages 46–53 to define its composition and analyze its relationship to other kidney types. By integrating single-cell RNA sequencing with *in situ* hybridization, immunostaining, and functional uptake assays, we identify mesonephric nephron segments, progenitor populations, and the progressive transition from mesenchymal precursors to differentiated epithelial tubules. Transcriptomic profiling of 14,411 cells reveals conserved nephron segment identities and shared differentiation programs revealed conserved nephron segment identities and a clear transition from mesenchymal progenitors to epithelialized tubules. Comparative analyses with published *Xenopus* pronephric and mammalian metanephric datasets identified strong transcriptional conservation between *Xenopus* mesonephric and mammalian metanephric nephrons. Functional assays confirmed the timing of filtrate uptake, marking the onset of renal function during metamorphosis. Together, these findings provide the first comprehensive single-cell map of *Xenopus* mesonephric development, demonstrating that the mechanisms of nephron patterning and differentiation are conserved across vertebrate kidney forms. This work establishes the *Xenopus* mesonephros as a robust model for studying vertebrate kidney development and evolutionary transitions among nephron types.

## Introduction

The kidney is an organ that functions to secrete soluble waste products from an organism. It performs this function through epithelial tubes called nephrons. The specialized tubules that make up each nephron take in fluid from the coelomic cavity or the vasculature, reabsorb solutes required by the organism, such as proteins, carbohydrates, and salts, and excrete waste products as urine. In freshwater organisms, nephrons remove the excess water, whereas in terrestrial animals, nephrons reabsorb water and therefore concentrate waste. Because nephron function depends on the precise patterning of solute carriers and ion channels, distinct genes are expressed within specific regions along the proximo-distal axis.^1–3^ Several kidney diseases preferentially manifest within specific nephron segments, such as certain cystic kidney diseases and glomerulopathies, highlighting differences in the development and function of these distinct nephron regions.^4–7^

The mammalian kidney develops in three consecutive forms, each exhibiting an increasingly complex arrangement of nephrons. This three stages are referred to as the pronephros, mesonephros and the metanephros.^8,9^ The first kidney form is the pronephros,^10^ which consists of a single nephron that originates in the cervical region of the intermediate mesoderm.^11^ In fish and amphibians, this structure is functional and performs many of the functions of the nephrons in later kidney forms.^12,13^ The most posterior region of the pronephros gives rise to the nephric duct, which is retained throughout development and provides the structural foundation for subsequent kidney forms. The second kidney form is the mesonephros, which develops in the caudal intermediate mesoderm and is composed of multiple nephrons that converge onto the nephric duct.^14^ This form contains true glomeruli connected to the vasculature.^15^ In fish and amphibians, the mesonephros represents the final adult kidney form, whereas in mammals it is remodeled into the Wolffian duct of the male reproductive tract.^16^ In amniotes, a third kidney stage, the metanephros, develops. This structure forms in the sacral region of the intermediate mesoderm and is initiated by branching of the nephric duct.^17,18^ Although each of these kidney forms is known to perform similar functions and arises through epithelization from mesenchymal progenitors, the extent to which cell types are conserved across kidney stages remains largely unexplored.

While the morphologic stages of mesonephric development have been described in many vertebrates, the precise molecular mechanisms and signaling pathways that drive mesonephric nephrogenesis remain less thoroughly investigated compared to the permanent metanephric kidney.^14,19^ In *Xenopus* and zebrafish, the pronephros is the preferred kidney form for developmental and disease studies because it develops rapidly, is functional, and can be directly visualized through the epidermis. This form has been widely used to model many diseases observed in the mammalian metanephric kidney.^20–22^ In mammals, the metanephric kidney is the preferred structure to study, as it most closely models human kidney disease and can be examined after birth.^23,24^ As a result, the mesonephric kidney has been largely overlooked. Much of our current understanding of the mesonephros is derived from structural observations of fully formed mesonephric kidneys in aquatic organisms or from indirect observations made during studies of early metanephric development in mammals.^15,25–28^ Moreover, most insights into mesonephric gene expression are based on a limited set of genes examined in zebrafish and mouse.^29,30^

It has been hypothesized that each successive form of the kidney reuses a shared genetic program to generate nephrons with similar cellular composition. This idea is supported by multiple studies showing that many phenotypes observed in mouse or human metanephric nephrons are recapitulated in the frog and zebrafish pronephros.^31^ In addition, single-cell sequencing (scRNA-seq) and *in situ* analysis of the functional *Xenopus* pronephros have reveled a conserved genetic profile relative to the adult mouse metanephric kidney.^12,32,33^ However, because these comparisons were performed across different species and kidney forms, the degree of conservation is difficult to quantify. To date, direct comparisons of cell types across distinct kidney forms within a single developmental framework have not been performed.

The goal of this study was to characterize the cell types and transcriptional programs that define the developing and mature mesonephric kidney in *Xenopus laevis*. To accomplish this, we used *X. laevis* tadpoles to perform a comparative analysis combining transgenic reporter lines, immunostaining, *in situ* hybridization, and single-cell RNA sequencing. These approaches enabled the identification and classification of nephron progenitors and differentiated cell types throughout mesonephric development. We found that early mesonephric nephrons contain distinct epithelial populations that progressively mature into conserved nephron segments. Transcriptomic comparisons revealed strong conservation of cell-type–specific gene expression between *Xenopus* mesonephric and mammalian metanephric kidneys. Together, these findings establish a molecular and cellular framework for mesonephric kidney development and highlight conserved pathways of nephron differentiation across vertebrates.

## Materials And Methods

### Xenopus laevis care

*Xenopus laevis* embryos were used as a vertebrate kidney model. Sex differentiation does not occur at the embryonic stages analyzed. Wild type *X. laevis* adult males and females were purchased from Nasco (LM00713M and LM00531MX). Transgenic *pax8*::GFP^< NXR_0132>^ cryo-sperm was obtained from the NXR^34,35^ (National *Xenopus* Resource) and was used via standard protocols ^34,35^. Adult *Xenopus laevis* were maintained under standard conditions in an AAALAC-accredited facility. Embryos were obtained by *in vitro* fertilization, staged according to Nieuwkoop and Faber, and maintained in maintained in recirculating aquatic housing at 18–20°C with 12-h light/dark cycle. *Xenopus* were induced to lay eggs using HCG and raised under normal conditions.^36,37^ All embryos were grown in 1/3X MMR (33 mM NaCl, 0.66mM KCl, 0.33 mM MgSO_4_, 0.66 mM CaCl_2_, 1.66 mM HEPES pH7.4). At around stage 45-47 embryos were fed a mixture of Spirulina (Nutricost) and macerated Frog Brittle (Nasco SA05960) dissolved in 1/3xMMR. All animal procedures were approved by the UTHealth’s Center for Laboratory Animal Medicine Animal Welfare Committee (protocol #: AWC-19-0081), which serves as the Institutions Animal Care and Use Committee (IACUC), and the IACUC at Icahn School of Medicine at Mount Sinai (protocol #16-1152 AR202100046 Yr2). To identify stages past stage 47 the formation of the limb was used. For all experiments, embryos were anesthetized in 0.02% Benzocaine (Spectrum BE130) diluted in 1xMMR (100mM NaCl, 2mM KCl, 1mMMgSO4, 2mM CaCl2, 5mM HEPES pH7.4) before fixation or manipulation, and all efforts were made to minimize animal use and suffering.

### Kidney isolation and dissociation of kidney cells

Wild type *X. laevis* were grown to stage 50 using standard protocols^38^. Manipulations were carried out in petri dishes containing 2% agar melted in 1xMMR. Kidneys were moved using a glass pipette and stored in a 1.5 ml tube in 1xMMR on ice until dissociation. Approximately 100 kidneys were dissected for further processing. Dissociations were carried out at room temperature unless otherwise noted. Kidneys were washed twice in 1xMMR and allowed to settle by gravity. Liquid was removed and replaced with 10mg/ml Collagenase Type IV for 30 min at room temperature until tissues were mostly disassociated. Cells were pelleted by centrifugation at 200xg for 3min, then the supernatant was replaced with 50:50 (1xTrypLE (Gibco 12605010):1xMMR) supplemented with 1mM L-cystine (Alpha Aesar A10435) + 10U/ml Papain (Sigma P4762) until a single cell suspension was obtained (∼1.5hrs). As constant shaking resulted in cell death, the tube was gently inverted once every 10 min and allowed to incubate on its side. Cells were strained threw a 40um sieve (VWR 732-2757). The sieve was washed with 25ml 45:45:10 1xMMR:L15:FCS. Cells were pelleted 500xg for 5min. Supernatant was removed and ∼500ul 1xMMR was added to the tube and cells were gently resuspended by shaking, then moved to a 1.5ml tube containing 1:1 1xMMR:Optiprep (Cosmo Bio Usa Inc AXS1114542) with a glass pipette. Cells were centrifuged 1000xg for 5 min and the upper and interphase layer was moved to a new tube with a glass pipette, leaving yolk pellets and cell debris in the denser lower layer. Cells were washed 2x with PBS by centrifugation 500xg 5 min to remove residual Optiprep and to prevent clumping.

### Immunostaining and in situ analysis

*In situ* analysis and antibody staining was performed using previously established protocols. ^39,40^ For immunostaining, embryos were washed with PBS and bleached in PBS + 0.6% H2O2 overnight at room temperature. Samples were blocked in 10% Goat Serum in P-triton (PBS + 0.01% triton-X100) for at least one hour. Antibodies/lectins 1:1000 αAc-tubulin (Sigma T6793), 1:1000 EC-lectin (Vector labs FL-1141-5) and 1:100 LEL-Lectin (vector labs DL-1178-1) were incubated overnight at 4°C. Imaging was taken on Zeiss LSM 800 confocal microscope. For *in situ* analysis, we used digoxigenin-labeled RNA probes that were prepared using a DIG RNA labeling kit (Roche). Constructs were linearized and synthesized using the listed enzyme and polymerase: *atp1a1*(T7, SmaI), *clcnkb* (T7, EcoRI), *foxi1* (T7, HindIII), *irx1* (T7, EcoRI), *irx3* (SP6, NcoI), *mecom* (SP6, SacII), *nphs1* (T7, SmaI)*, slc4a4* (T7, SalI)*, slc5a1* (T7, SmaI)*, six1* (StuI, T7)*, lhx1* (T7, XhoI)^12,32,41–43^. At least 5 embryos by condition were processed as described, eliminating the RNAse A/T1 step^44^. 1:3000 dilution of Anti-DIG Fab fragments (Roche) and NBT/BCIP tablets (Roche) were used to detect probes. A *six1* in situ probe was generated by cloning full length cDNA into pCS2-6xMYC. *six1* was PCR amplified from stage 30 cDNA using primers Six1F-ttcgAGGCCTatgtctatgctgccttcctttggc and Six1R-tagtTCTAGAttacgatcccagatccaccaggctggagg. Product was digested with StuI or XbaI and cloned into the same sites of pCS2-6xMYC. Plasmid was linearized with StuI therefore the Myc tag was not included in the probe.

### Single-cell mRNA sequencing (scRNA-seq)

scRNA-seq was performed at UTHealth Cancer Genomics Center (CGC) and processed using Seurat^45–47^. Quality control, data processing and statistics are founded in the Supplementary information file.

### Study design and experimental units

#### Groups being compared, including controls

This study was designed as a comparative descriptive and transcriptomic analysis rather than a treatment-based experiment. Therefore, no experimental treatment or control groups were applied. The main comparisons were developmental (between early and late mesonephric stages) and cross-species (between *Xenopus laevis*, and mammalian datasets). Except for the functional assay – EC-Lectin and Hoechst dye injection, where the pronephros staining was used as a positive control. In all staining experiments we included appropriate positive and negative controls to ensure specificity and reproducibility. In *in situ* hybridization assays, sense probes were used as negative controls to confirm the absence of nonspecific staining, and colorimetric reactions were stopped when the sense control showed background staining.

#### Experimental unit

The experimental unit was the “individual embryo” for *in situ* hybridization and imaging experiments, and the individual dissected mesonephric kidney for single-cell RNA sequencing analyses. For validation imaging and gene expression comparison across datasets, each condition was confirmed in at least three biological replicates, representing independent embryo clutches. The total number of animals used across all experiments was approximately 300 *Xenopus laevis* tadpoles in total for this study and only morphologically normal *Xenopus laevis* embryos at the appropriate NF stages were included in all experiments.

#### Sample size justification

No formal *a priori* statistical power calculation was performed, as the study was primarily descriptive and exploratory, aimed at defining mesonephric cell types and developmental trajectories. Sample sizes were determined based on established standards from prior *Xenopus* kidney studies, which demonstrated reproducible cellular resolution using similar embryo numbers^21,31,33,48,49^. Sample numbers were empirically optimized to 1) ensure sufficient tissue for single-cell RNA sequencing library generation (minimum ∼10,000 cells per dataset), 2) achieve clear and consistent staining and imaging quality at each developmental stage and balance biological reproducibility with the ethical principles of animal use (Replacement, Reduction, and Refinement).

## Results

### Evaluating the timing of mesonephric development

To visualize the onset of mesonephric development, we analyzed *pax8::GFP* transgenic animals (RRID:NXR_0132)^50^ (Fig. 1A). Because external morphology can vary between individuals, developmental stage was initially assessed using limb bud morphology. However, we observed that mesonephric development showed variability even among animals with comparable limb bud staging, indicating that external morphological criteria alone do not precisely predict kidney maturation.

**Figure 1:**
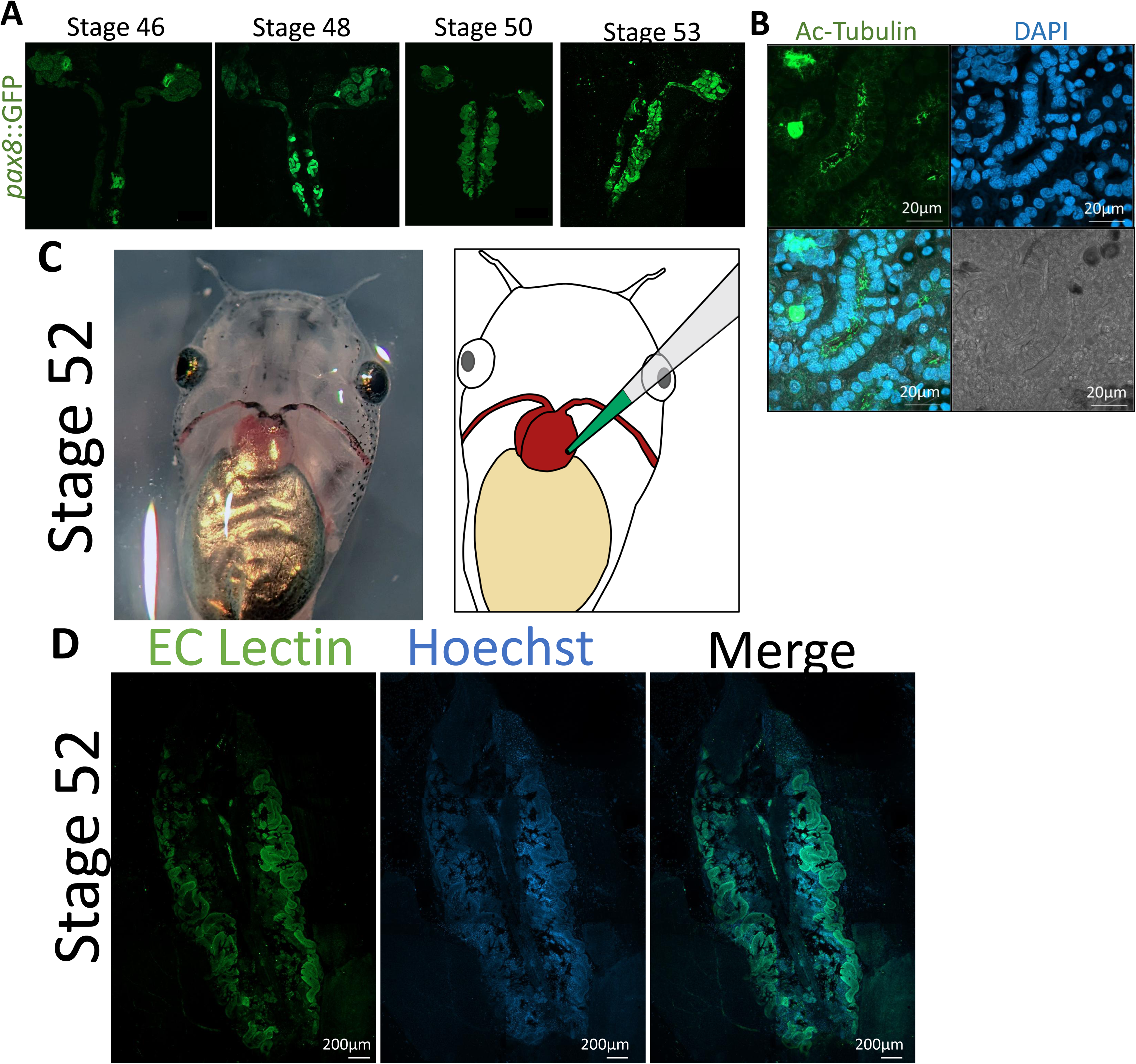
Visualization of mesonephric kidney development and function. A) Live imaging of *pax8*::GFP embryos over the stages of mesonephric development. B) Paraffin embedded section of adult mesonephric kidney tubules stained with acetylated tubulin (green), DAPI (blue) and Differential Interference Contrast (DIC) image. C) Ventral side of stage 52 embryo along with illustration showing injection of Hoechst and EC-lectin (green) into the heart (brown). D) Image of the mesonephric kidney after injection of EC-lectin (green) and Hoechst 3342 (blue) into the heart.

In *Xenopus*, both pax8::GFP and *pax8 in situ* hybridization probes are well-established markers of pronephric nephrons.^50^ *pax8* is expressed in kidney epithelial cells as well as nephron progenitor populations in the *Xenopus* pronephros and the mammalian metanephros.^51–53^ We used the classical descriptions of *Xenopus* development by Nieuwkoop and Faber (NF, 1956) to identify the different stages of mesonephros development and we detected the earliest signs of mesonephric nephron formation around stage 46. Notably, the first mesonephric nephrons formed near the posterior end of the nephric duct and subsequently appeared progressively along its length. After stage 53, the identification of newly forming nephrons became more challenging due to the increased number of nephrons and the accumulation of melanocytes that obscure visualization. As expected, newly formed mesonephric nephrons displayed a ciliated cuboidal epithelial morphology (Fig. 1B), consistent with previous histological descriptions^54^.

To determine when the mesonephric kidney becomes functional, we adapted a dye-injection assay originally described by Zhao and Vize (2004).^12,48,55^ In this assay, labeled molecules are introduced into the circulation and their passage through the nephron is monitored. Because the mesonephric kidney cannot be readily visualized in live animals, we selected stable fluorescent tracers to assess filtrate entry into nephron lumens. A mixture of Hoechst 3342 and EC-Lectin (Erythrina cristagalli lectin) was injected into the cardiac ventricle of *X. laevis* tadpoles (Fig. 1C,D). Hoechst served as an injection control, as it labels nuclei in tissues reached by the circulation, confirming successful systemic delivery.

EC-lectin is a non-cell permeable carbohydrate-binding reagent that labels the apical surface of the proximal tubules cells when present within the nephron lumen. In permeabilized tissue, lectins can directly bind to both cell-surface and intracellular epithelial glycoconjugates (glycoproteins and glycolipids); however, in non-permeabilized animals, luminal staining is only observed when circulating lectin enters the nephron lumen. Therefore, detection of luminal EC-lectin signal indicates that filtrate has entered the nephron. Because dye entry into the mesonephros could theoretically occur via glomerular filtration or via nephrostomal connections to the coelomic cavity, we interpret lectin staining as evidence of functional luminal flow rather than as a strict measure of glomerular filtration alone.

Lectin labeling intensity was variable between animals, and some injections failed to produce detectable staining in either kidney form. The pronephros was used as an internal positive control; mesonephric lectin staining was only observed in animals in which pronephric staining was also detected. Mesonephric luminal staining was first consistently observed around stage 50, suggesting that the earliest mesonephric nephrons become functionally integrated into the excretory system at this stage. This timing is consistent with NF descriptions of the onset of mesonephric nephron maturation.

### Transcriptome analysis of the mesonephros

To define the cellular composition of the *Xenopus* mesonephric kidney, single-cell mRNA sequencing analysis was performed. Kidneys from stage 50-52 were selected because nephrons are functional at these stages, yet the organ continues to grow and retain progenitor populations. In addition, stages 52-56 were included because the kidney remains sufficiently thin for whole-mount staining, facilitating parallel anatomical validation, and multiple biological replicates could be obtained within a relatively short timeframe (∼3 weeks) (Fig 1 C, D). At these stages, the kidney is readily identifiable without molecular markers, as pigment cells outline the connective tissue covering the ventral surface (Fig. 2A).

**Figure 2:**
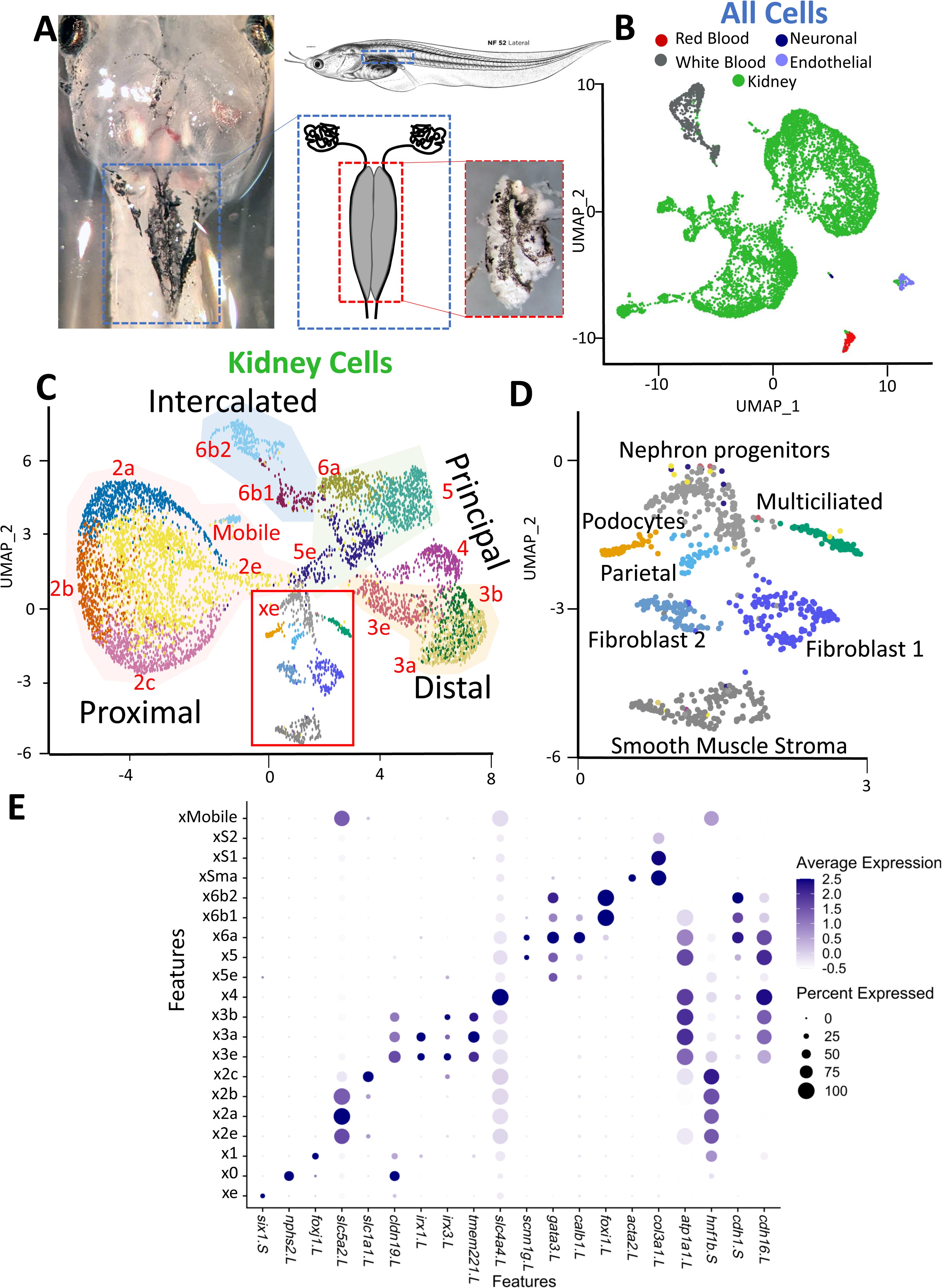
Single-cell mRNA sequencing of stage 49-52 mesonephric kidneys. A) Diagram of kidney isolation on stage 52 *X. laevis,* along with image of kidney before and after isolation. Blue dotted lines surround both pronephric and mesonephric kidney. Orange dotted lines indicate region of kidney isolated. B) UMAP showing cell types isolated. C-D) Analysis of subclustered kidney cells. C) UMAP of kidney cell types. D) Magnified image of orange box in panel C. E) Dotplot showing expression of markers in the mesonephros that have been shown to define each segment in the pronephros.

A total of 100 mesonephric kidneys were manually dissected from heavily anesthetized animals (100 µg/ml benzocaine). Upon dissection, the kidneys contracted along the anterior–posterior axis, suggesting that the organ is maintained under mechanical tension *in vivo*. Following dissociation, approximately 1 × 10^7^ cells were obtained. After sequencing, stringent quality control filtering was applied. Cells with low read counts were excluded to minimize inclusion of poorly sequenced cells or cellular fragments. After filtering, 14,411 high-quality cells remained, with an average sequencing depth of ∼34,500 reads per cell.

Because nephron segment nomenclature in *Xenopus* evolved over time, we adopted, where possible, the segmentation framework described by Corkins et al. (2023) for the pronephric nephron to maintain consistency across datasets. Clustering and UMAP embedding were performed independently from the previously published pronephric scRNA-seq dataset to avoid bias in cluster assignment.^33^

To restrict downstream analyses to renal populations, non-kidney cells were separated prior to detailed subclustering. Canonical nephric markers from the *Xenopus* pronephros and mouse metanephros were used to define kidney identity. Mesonephric cells were identified by expression of *cdh1*, *cdh16*, *hnf1b*, *atp1a1*, *nphs2*, and *foxi1* and subclustered away from non-kidney cells such as red blood cells (Fig. 2B,C: Fig. S1). Clusters associated with nephric populations in UMAP space and expressing these markers were retained in the kidney subset. Excluded populations included three white blood cell types, two endothelial populations, and red blood cells; all other clusters exhibited transcriptional association with the nephric compartment (Fig. S2).

We identified many of the same epithelial cell types previously described in the pronephric single-cell dataset (Fig. 2C). To facilitate comparison, these were numbered 1–6 to match the pronephric classification. Broadly, these populations fall into proximal distal and duct categories, with duct cells comprising both principal and intercalated cell types (Fig. 2C). In addition to these conserved epithelial populations, mesonephros-specific cell types were identified, including an immature parietal epithelial population and three stromal subtypes. The stromal populations consisted of two extracellular matrix–producing fibroblast-like groups and one smooth muscle–like group (Fig. 2D).

Nephron epithelial clusters were connected by intermediate populations, including a central group of cells expressing nephron progenitor-associated markers such as *six1*. These progenitor-like populations were labeled with an “e” designation (e.g. 2e, 3e, 5e) to denote embryonic or early states (Fig. 2C,E). The presence of these transitional clusters suggests that the mesonephros contains a continuum of differentiation states, consistent with ongoing nephrogenesis at these developmental stages.

### Comparative transcriptomics of Xenopus nephron forms

To determine the similarities and differences between the *Xenopus* kidney stages, pronephric and mesonephric single-cell datasets were integrated (Fig. 3). To ensure comparability, the pronephric dataset was reprocessed using the updated *X. laevis* 10.1 genome assembly prior to integration. This minimized annotation bias and enabled direct transcript-level comparison across developmental stages. Overall, clustering revealed strong correspondence between nephron segments across the two stages. Except for previously described cell types, the most notable divergence was observed in the distal cluster x6a from the pronephros, which did not co-cluster with its mesonephric counterpart (Fig. 3A). These cells represent the most distal region of the duct. In the pronephros, this population likely derives from cloacal or adjacent epidermal cells, whereas in the mesonephros, this area appears to originate from earlier-stage distal principal-like cells. The distinct clustering of x6a therefore likely reflects differences in developmental origin and tissue context rather than complete functional divergence.

**Figure 3:**
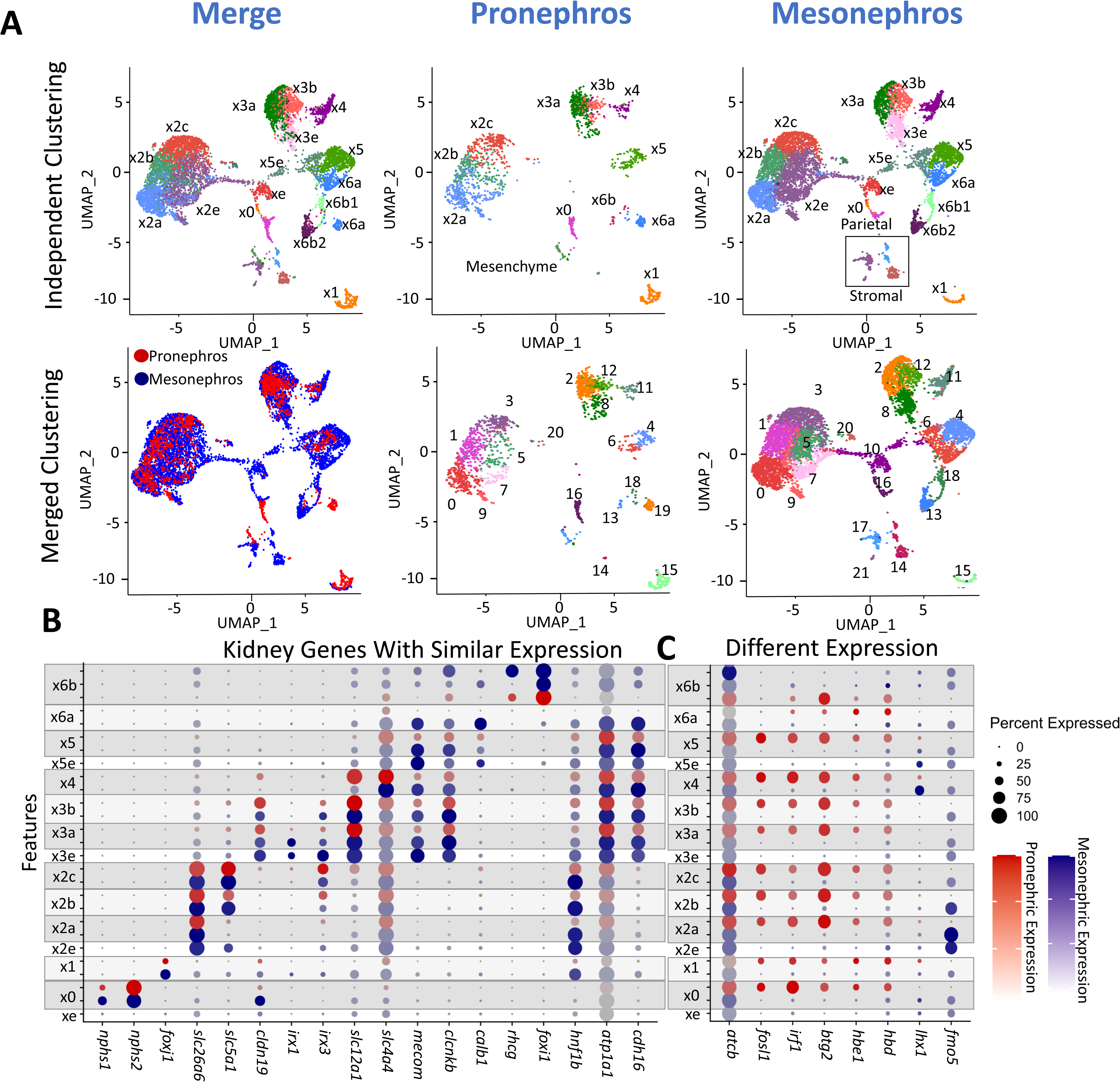
Integration of single-cell data between mesonephric and pronephric kidneys. A) UMAP showing the integrated dataset. Cluster names as identified by independent analysis, along with separated pronephros and mesonephric samples. Clustering was performed on the joint sample to indicate similar cell types. B) Dotplot showing normalized expression of kidney gene markers for each segment. Orange dots show expression in the pronephros and blue dots indicate expression in the mesonephros. C) Dotplot showing genes that have different expression between the pronephros and mesonephros.

The mesonephric dataset contained a larger number of intercalated cells, allowing more detailed resolution of this population. These cells segregated into two transcriptionally distinct groups (x6b1 and x6b2) that correlate with type A and type B intercalated cells described in mammalian kidneys (Fig. 3B). Marker expression patterns were consistent with these identities, supporting conservation of ionocyte specialization across vertebrates. In addition, a small cluster expressed extracellular proteases including *mpo*, *mmp7*, *mmp9*, *prtn3* and *prss57*. These genes are commonly associated with motile or infiltrating cell populations. Based on this signature, we provisionally labeled this cluster as “Mobile”. Notably, these cells also expressed markers characteristic of cluster x2a, corresponding to the most proximal tubule population (Fig. 2E). This overlap may indicate transitional or remodeling states during nephron expansion. However, because protease expression can also reflect tissue remodeling or immune-associated processes, we interpret this population conservatively as a transient or migratory-like cell state rather than definitively migratory proximal tubule cells.

Within individual nephron segments, surprisingly few transcriptional differences were detected between pronephric and mesonephric counterparts. Genes such as *irf1*, *btg2*, *fosl1* and *jun* were more highly expressed in the pronephros (Fig. 3C). These genes are associated with stress response and early development activation, and their enrichment likely reflects differences in nephron maturity and/or increased dissociation sensitivity in earlier stage tissue. Additional differences included expression of hemoglobin genes (*hbe1* and *hbd*) in pronephric tubules, though the biological basis for this remains unclear and may reflect transient developmental or technical influences.

One notable gene showing differential retention was *lhx1* (Fig. 3C). In the pronephros *lhx1* is required for nephron formation and is ubiquitously expressed in the nephron prior to epithelization, later becoming restricted to the nephrostome and migrating duct cells after epithelization.^41,56^ In contrast, *lhx1* expression persists in mesonephric principal cluster 4. A similar pattern is observed in mouse, where it is initially broad and later becomes restricted to distal convoluted tubule segments. Except for the *lhx1* expression, transcriptional signatures of nephron segments were highly conserved between pronephric and mesonephric kidneys, supporting strong molecular continuity across developmental stages.

### Mesonephric progenitor lineage

Because the mesonephric dataset captures a continuum of nephron maturation, we performed trajectory analysis to establish lineage relationships (Fig. 3 A-C). Less differentiated cells were identified by expression of early kidney markers including *wt1*, *six1, meis1*, and *lhx1.* These cells were labeled with the “e” (embryonic/early) label (Fig 2C). In contrast, more differentiated populations expressed higher levels of mature epithelial markers, including transporters and junctional components (Fig. 2E), and co-clustered with mature pronephric cells in the integrated analysis (Fig. 3A). To further distinguish proliferative from more mature states, we analyzed cell cycle signatures using published S-phase and G2/M gene sets commonly applied in mouse and human single-cell studies.^57^ Cells were assigned cell cycle phase based on enrichment of these gene sets. Early populations showed higher proportions of cycling cells, whereas more differentiated clusters were enriched for G1/G0 signatures, consistent with reduced proliferation accompanying nephron maturation.

To infer developmental trajectories, we applied Monocle 3 for pseudotime analysis (Fig. 4A-D).^58^ Early, intermediate and late gene expression dynamics were validated using lineage specific markers (Fig. 4 E-G). While trajectory topology varied modestly depending on UMAP embedding and clustering parameters in Seurat, several consistent features emerged across analyses. Specifically, four principal branches arose from the early progenitor population: 1) A stromal and multiciliated lineage, 2) A podocyte and parietal epithelial lineage, 3) A proximal tubule lineage, 4) A distal nephron lineage, including intercalated, principal, and distal cell types (Fig. 4B). The separation of podocyte, stromal, and multiciliated lineages was more clearly resolved when subclustered independently from fully differentiated epithelial populations (Fig 4C-D), indicating that inclusion of mature cell types can obscure early branching structure. These four developmental trajectories parallel those described in mammalian mesonephric kidneys and exhibit similar lineage-associated gene expression patterns, suggesting conservation of nephron patterning programs. Collectively, these data indicate that mesonephric nephron progenitor cells generate three primary nephrogenic axes: proximal epithelial, distal/principal/ionocyte, and stromal/glomerular lineages, which together establish the structural and functional organization of the mature nephron.

**Figure 4:**
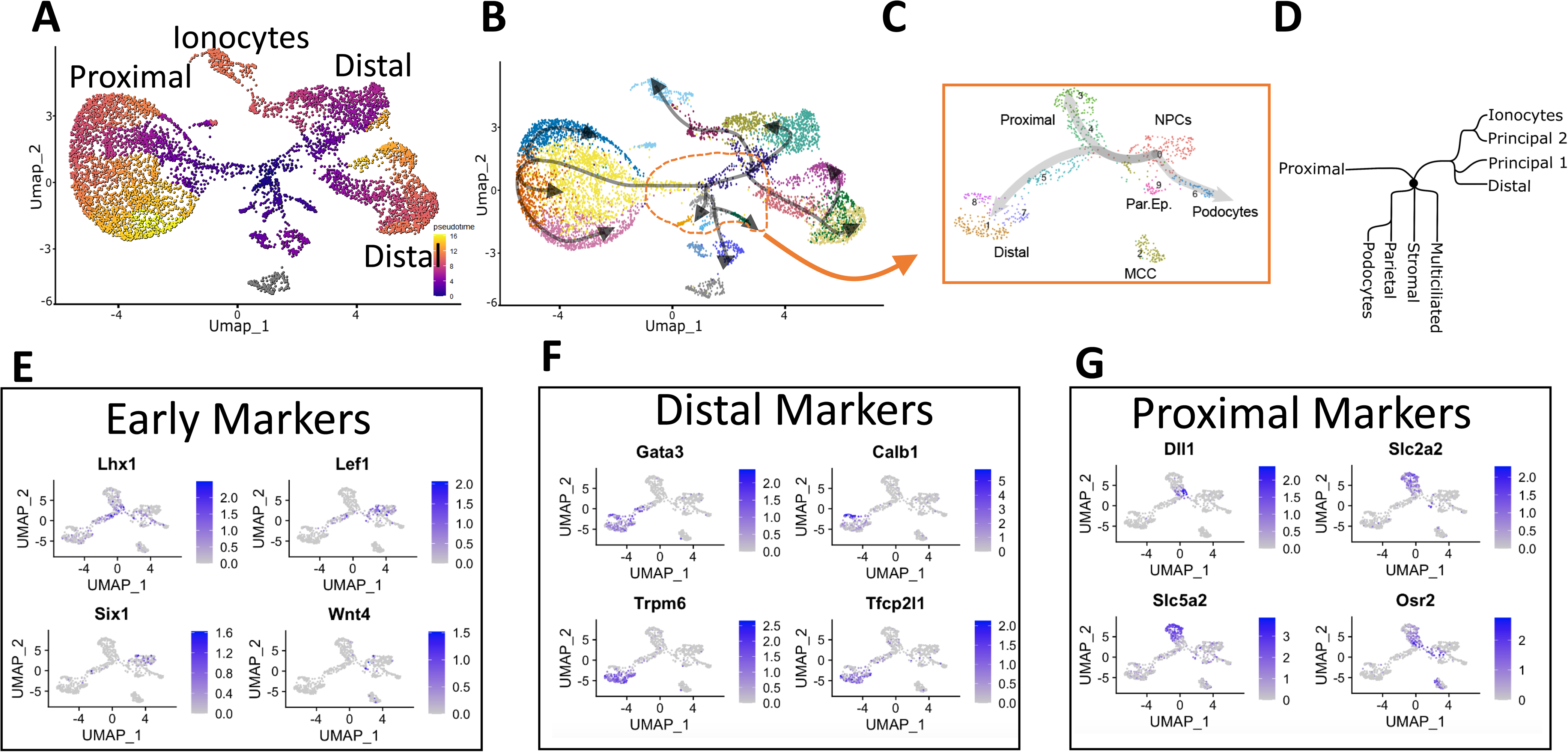
Pseudotime analysis of the developing *Xenopus* mesonephric kidney. A-B) Pseudotime trajectory analysis was performed using Monocle 3. Darker blue colors indicate more embryonic lineages, while brighter yellow indicates more differentiated cell states. Multiciliated cells were not analyzed (indicated in grey). Gray arrows indicate the predicted developmental trajectories inferred by Monocle 3. C) Subset of embryonic cell types indicated by orange dashed outline in B. D) Predicted lineage relationships among identified mesonephric cell types. E-G) UMAP plots showing the expression of marker genes associated with specific cell types within the early, less differentiated population.

### Evolutionary conservation of kidney development

To determine whether the developmental programs are conserved across vertebrates, we compared the *Xenopus* mesonephric single-cell dataset with published adult mouse metanephric kidney data (Fig. 5). Integration was performed using an approach analogous to that used for the pronephros–mesonephros comparison, enabling cross-species alignment of transcriptionally homologous cell states.

**Figure 5:**
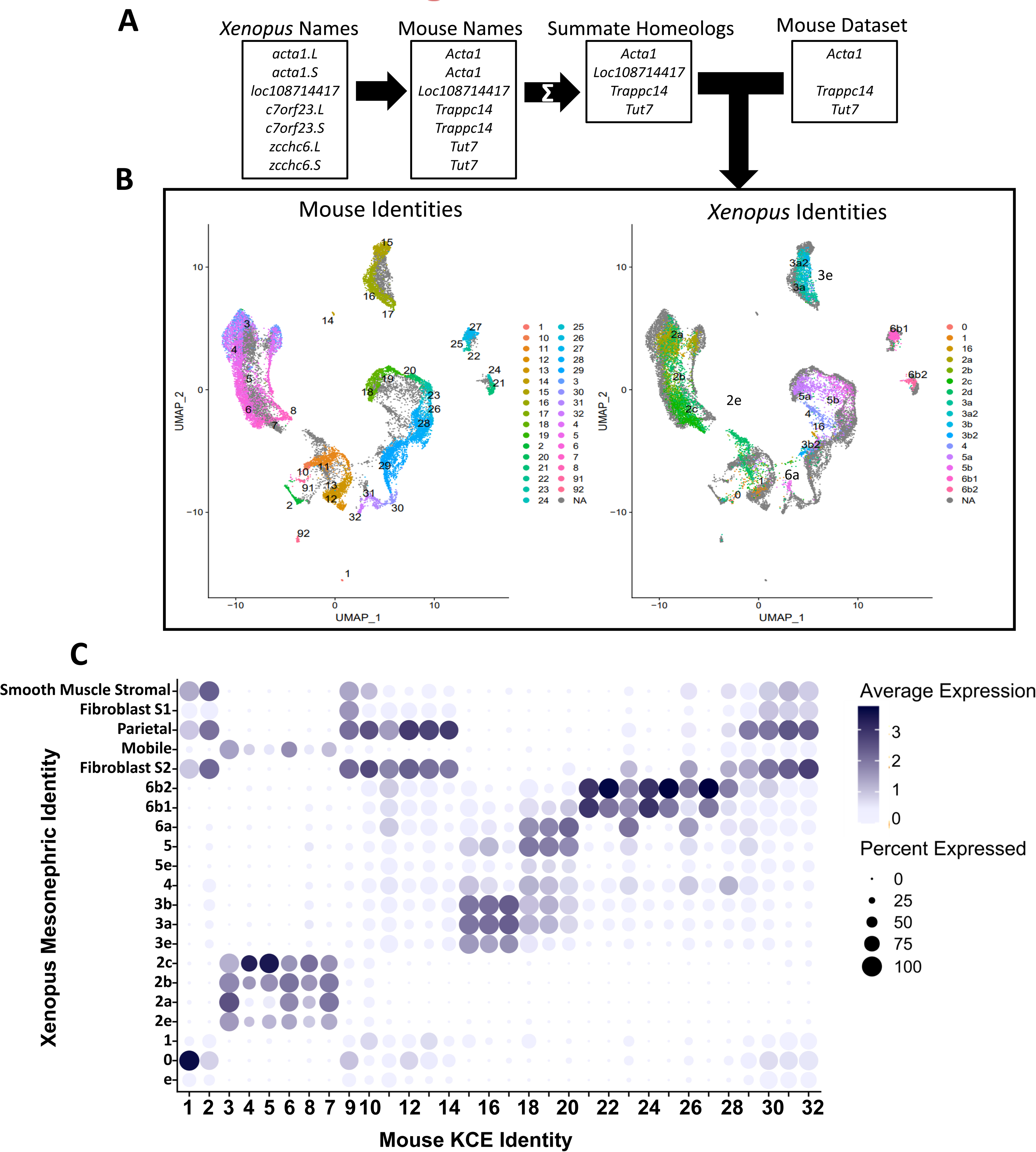
Comparison between the cell types of the *Xenopus* mesonephric kidney and the adult mouse metanephric kidney. A) Methodology to convert frog data to be compatible with mouse and human datasets. B) UMAP projection showing integrated mouse metanephric and *Xenopus* mesonephric data sets. C) Using the Mouse Kidney Cell Explorer dataset [1], the top 50 enriched genes for each cluster were identified using the FindAllMarkers function in Seurat. Using the AddModuleScore [2] function average relative expression for each gene set was calculated for the *Xenopus* Mesonephric Kidney. Average expression, and percentage of cells that have at least one transcript was displayed for each cluster. Genes with no clear homeolog were ignored. 1. Ransick A, Lindstrom NO, Liu J, et al. Single-Cell Profiling Reveals Sex, Lineage, and Regional Diversity in the Mouse Kidney. Dev Cell 2019; 51: 399-413 e397. 2. Tirosh I, Izar B, Prakadan SM, et al. Dissecting the multicellular ecosystem of metastatic melanoma by single-cell RNA-seq. Science 2016; 352: 189-196.

Because the adult mouse metanephros is fully differentiated, it contains relatively few undifferentiated or progenitor-like populations compared to the *Xenopus* mesonephros, which includes nephrons at multiple developmental stages. As expected from the strong correspondence observed between the pronephros and mesonephros we detected substantial conservation between mature mesonephric epithelial populations and mouse adult nephron segments (Fig. 5B). Proximal tubule, distal tubule, principal, and intercalated populations co-clustered across species based on shared transporter expression profiles and segment-specific transcription factors, supporting conservation of terminal epithelial identities (Fig. 5C).

Consistent with our earlier observations, *Xenopus* cell types lacking clear one-to-one mammalian analogs did not align with discrete mouse epithelial segments. Instead, multiciliated cells and nephron progenitor–like populations grouped transcriptionally near Loop of Henle (LOH) clusters in the integrated embedding. Notably, amphibians lack a canonical Loop of Henle structure, and therefore this association is likely to reflect partial overlap in shared gene modules rather than strict anatomical equivalence. For example, enrichment of solute transporter programs and epithelial polarity genes may drive proximity in integrated space without implying direct homology of segment architecture.

Importantly, early mesonephric progenitor populations did not form distinct cross-species clusters with adult mouse nephron progenitors, consistent with the developmental maturity of the mouse dataset. Rather than representing conserved progenitor states *per se*, cross-species alignment was strongest among differentiated epithelial lineages. This pattern suggests that terminal nephron segment identities are more transcriptionally conserved than transient developmental intermediates when comparing across species and developmental stages.

Taken together, these results support strong evolutionary conservation of core nephron epithelial programs between amphibian mesonephric and mammalian metanephric kidneys, while also highlighting lineage- and stage-specific differences, particularly in progenitor and amphibian-specific multiciliated populations.

### Spatial positioning of cell types within the mesonephros

To determine the anatomical localization of the transcriptionally defined cell populations within the *Xenopus* mesonephros, we examined spatial expression patterns of established kidney markers. We first set out to identify the location of the kidney progenitor cells. In the pronephros, *pax8* and *lhx1* are commonly used to label early nephric progenitors. Although *pax8* marks nephron progenitors in both the pronephros and mammalian metanephros, expression of the *pax8*::GFP transgene in the mesonephros was restricted to epithelialized tubules (Fig 1A), suggesting that in this context *pax8* labels differentiated epithelial cells rather than early progenitors. Unlike *pax8,* which is constitutively expressed in the entire nephron, *lhx1* is only present in the pronephric kidney mesenchyme, with its expression decreasing shortly after epithelization. However, neither Lhx1 protein nor *lhx1* mRNA was detected using antibody staining or *in situ* analysis before stage 50.^41,56^ However, expression was readily observed in the brain, with only a weak nephric signal emerging around stage 52 embryos near the midline (Fig. S1B). This nephric signal corresponded spatially to the early principal cell population identified in cluster 4 by scRNA-seq, suggesting that *lhx1* expression at these stages reflects early epithelial differentiation rather than multipotent progenitors.

Based on the scRNA-seq analysis, *six1* emerged as a more specific marker of mesonephric progenitor cells and it has also been described as a transient early kidney progenitor marker in mouse.^59^ A *six1* probe was cloned from cDNA and used for whole-mount *in situ* hybridization. Strong staining was observed in collagen-rich cranial structures, and weaker signal was detected in the somatic myotome, consistent with known expression domains. Within the developing mesonephros, *six1* expression appeared as stripes of cells posterior to the nephric tubules at early stages (Fig. S1C). By stage 50, scattered *six1*-positive cells were distributed around the kidney, with enrichment near the dorsal midline (Fig 6A). In more developed kidneys (Stage 53), discrete clusters of *six1*-positive cells formed a linear arrangement along the dorsal side of the organ. This spatial pattern suggests that new nephron formation occurs adjacent to dorsal midline structures. The relatively weak signal and the limited number of progenitor-like cells identified by scRNA-seq indicate that these cells are rare and likely transient during mesonephric development.

**Figure 6:**
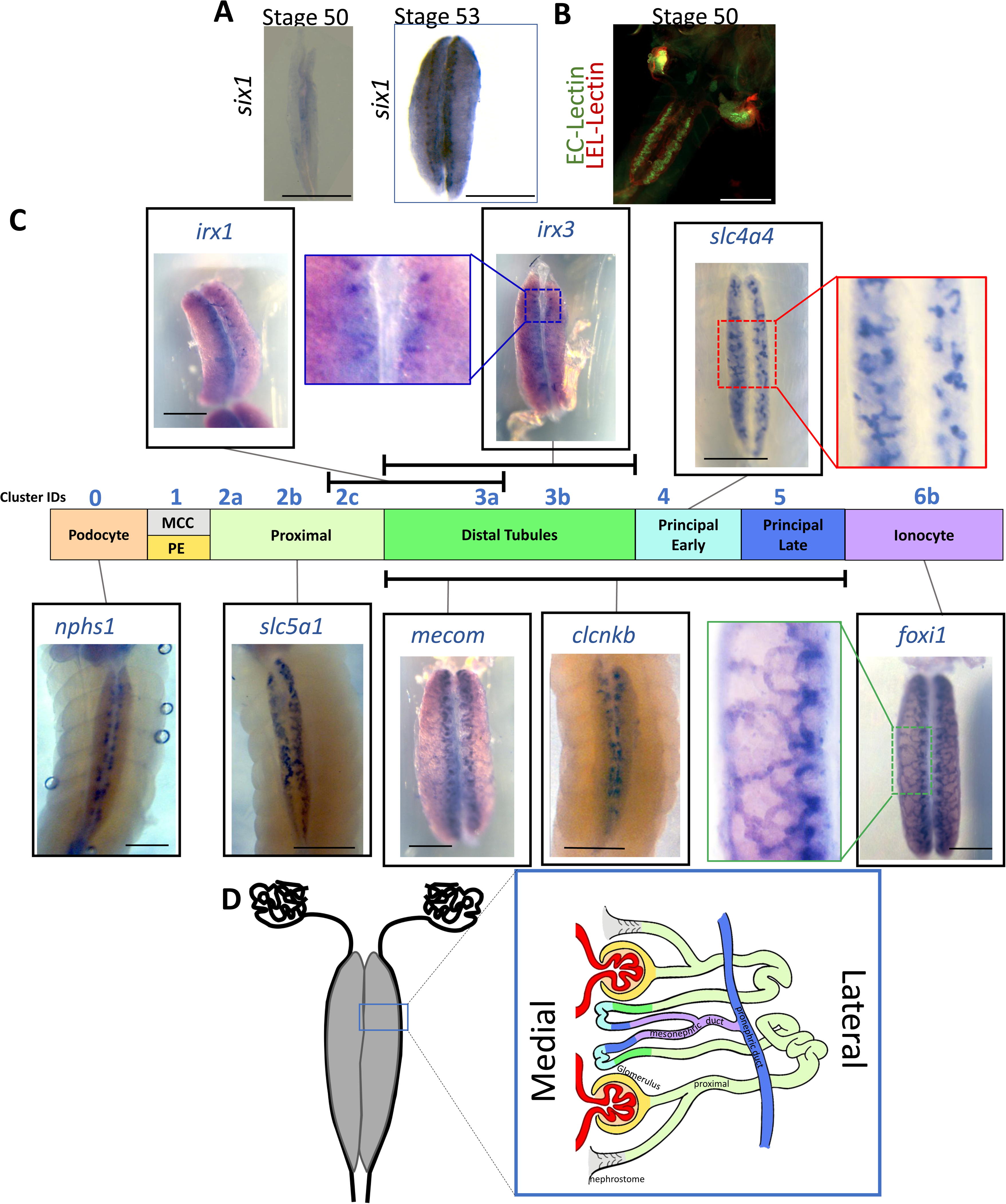
Spatial analysis of mesonephric cell types. A) *In situ* analysis of *six1* kidney progenitor marker B) Ec-Lectin::FITC and LEL-Lectin::Dylight594 staining of fixed stage 50 embryos. C) *In situ* analysis of nephron segments. Stage 52-53 kidneys were stained for indicated probes. Segments for each probe labels are indicated by the layout of the *Xenopus* nephron from proximal to distal. Blue numbers above each segment indicate clusters identified by single-cell analysis. Black or white scale bars indicate 1mM. D) Diagram indicating the model of mesonephric nephron structure.

To define the broader spatial organization of nephron segments, we first labeled proximal tubules using FITC-conjugated Erythrina cristagalli lectin (EC-lectin) (Fig 6B)^60^, which has been shown to selectively bind proximal tubular epithelium. Proximal tubules were predominantly located along the lateral aspects of the kidney and comprised the largest proportion of the organ, suggesting that most of the mesonephric nephron is composed of proximal tubules. This pattern was confirmed by *in situ* hybridization for *slc5a1,* a proximal tubule transporter (Fig 6C), consistent with both scRNA-seq data and prior reports.^32,49,61^

Podocytes were visualized using *nphs1*, which marks glomerular filtration barrier (glomerulus/glomus).^62^ *nphs1*-positive cells appeared as puncta structures along the medial regions of the kidney, defining the most proximal segment of the nephron. The positioning of these glomerular structures aligns with the renal artery, which runs along the midline between the paired kidneys, consistent with previous ultrastructural analyses of mesonephric vasculature.^15^

Distal tubule segments were identified using *irx1* and *irx3*.^63^ These markers labeled individual tubules along the medial edge of the kidney. *Mecom* (*evi1*) and *clcnkb* were used to label distal and principal cell populations.^64–66^ These genes strongly marked tubules positioned near the midline, with weaker projections extending dorsally toward the collecting duct. Although labeling intensity differed, the spatial distribution of principal cells was like that observed for *irx1/3-*positive distal segments.

Ionocytes/intercalated cells were identified using *foxi1*^67^, which labeled which labeled a series of converging tubules along the dorsal side of the kidney. This dorsal localization corresponds to the distal nephron and ductal regions identified transcriptionally in the scRNA-seq dataset.

Together, these data support a spatial organization in which nephrogenesis initiates medially and dorsally, as suggested by the distribution of *six1*-positive progenitor cells. Proximal tubules extend laterally and constitute most of the nephron mass (Fig 6D). These proximal segments connect medially to distal tubules positioned near the glomeruli. Distal segments then converge dorsally into the mesonephric duct, which ultimately connects to the pronephric duct for excretion. This integrated spatial map aligns transcriptional identities with anatomical position and provides structural context for the developmental trajectories inferred from single-cell analysis.

## Discussion

Relatively little is known about the molecular and developmental organization of the mesonephric kidney. Here, we combine single-cell transcriptomics with spatial validation to define the cellular composition, developmental trajectories, and evolutionary conservation of the *Xenopus laevis* mesonephros. Using existing single-cell datasets of the pronephros, we directly compare successive kidney developmental stages within the same vertebrate species.^33^ Our findings demonstrate that the mesonephros is composed of transcriptionally conserved nephron segments organized along proximal–distal axes that closely parallel both the amphibian pronephros and the mammalian metanephros, while also revealing stage-specific and amphibian-specific features.

### *Sequential* and asynchronous nephrogenesis

At stage 53, we observed approximately 20-30 glomeruli per kidney. In contrast, adult animals possess roughly 2000 nephrons, ^15,68^ indicating that only a small fraction of the final nephron number has formed at this stage. Posterior nephrons appeared to develop earlier than anterior nephrons, suggesting sequential nephron formation along the posterior–anterior axis. This spatial gradient indicates that mesonephric nephrogenesis proceeds asynchronously over an extended developmental window. Such prolonged and staggered nephron formation mirrors patterns observed in the mammalian metanephros. In humans, the first glomeruli form around week 9 of gestation, and nephron production proceeds for roughly 27 additional weeks, ultimately yielding approximately 250,000 to more than one million nephrons per kidney.^69^ The presence of asynchronous nephrogenesis in amphibian and mammalian kidneys suggests that iterative nephron addition over time represents a conserved developmental strategy rather than a lineage-specific adaptation.

### Cellular composition and tissue isolation

In contrast to our prior pronephric dataset, relatively few non-nephron cells were identified in the mesonephric single-cell analysis.^33^ This likely reflects the connective tissue capsule surrounding the mesonephros, which physically isolates the organ and reduces contamination during dissection. Endothelial and blood cell populations were nevertheless detected, as expected for a vascularized and functional kidney (Fig. S3). These cell types were expected, given their role in renal functional maturation.

Although early germline formation has been reported around stage 52, ^70,71^ and the mesonephros contributes to the male reproductive organ, we did not detect transcriptional signatures that clearly distinguished sex at the analyzed stages. Canonical sex-determining genes (*dmrt1, foxl2, dm-w*) and genes reported to show sexual dimorphism in mouse kidney did not segregate into sex-specific clusters. These genes were either completely absent or ubiquitous expressed in the proximal tubules. Sexual dimorphism may emerge later or may involve regulatory mechanisms not resolved at the transcriptional level at these developmental stages.^72^

### Conservation of nephron segment identity across kidney forms

A striking finding of this study is the strong transcriptional similarity between mature epithelial populations of the mesonephros and those of the pronephros. Proximal, distal, principal, and intercalated cell types exhibit conserved transporter repertoires and transcription factor expression profiles across kidney stages. These data suggest that successive kidney forms in *Xenopus* reuse a shared developmental program to generate segmented nephrons. Cross-species integration with adult mouse kidney further demonstrates that mature epithelial identities are deeply conserved across vertebrates. Despite anatomical differences between the amphibian mesonephros and the mammalian metanephros, terminal nephron segment identities cluster by shared functional gene modules rather than species of origin. This conservation implies that core epithelial patterning programs were established early in vertebrate evolution and have been redeployed in morphologically distinct kidney architectures.

### Temporal divergence of intermediate states

While mature epithelial populations are highly conserved, intermediate and progenitor-like states show greater divergence across species and developmental stages. Regulatory genes such as *osr1*, *dll1*, and *lhx1* are present in both amphibian and mammalian systems, yet their temporal differences in the gene expression differ. For example, DLL1 in human kidney development is broadly expressed across nephric lineages early and later becomes restricted to proximal segments^73^. In *Xenopus*, *dll1* expression corresponds more tightly with the divergence of proximal and distal lineages. A similar observation was made for *lef1*. This early divergence is likely driven by differences in the temporal regulation of gene expression between *Xenopus* and humans, which nonetheless converge on a broadly conserved developmental program for nephron formation.

### An amphibian-specific epithelial state

Cluster 4 (early principal cells) represents an unusual epithelial population that may reflect amphibian-specific physiological adaptation or developmental plasticity. Although its transcriptional profile aligns with principal cell identity in cross-species analysis, pseudotime reconstruction indicates that it develops from distal tubule populations rather than directly from the progenitors. Furthermore, the two most highly expressed transporters found in this population are *slc4a4* and *slc9a3,* which encode proteins necessary for pH balance within the proximal tubules in mammals,^74^ suggests partial integration of gene modules normally segregated into distinct nephron segments.

Spatial analysis does not fully resolve the anatomical positioning of this population, and *slc4a4* expression does not localize precisely with other distal or principal markers. Together, these observations suggest that amphibian mesonephric epithelial identities may exhibit greater transcriptional modularity than their mammalian counterparts. Rather than representing a simple positional mismatch, cluster 4 may illustrate flexibility in how conserved transport programs are combined to meet species-specific physiological demands. Further functional analysis will be required to determine whether this population reflects a transient developmental state or a stable amphibian-specific epithelial identity.

### Integrated spatial and transcriptional framework

By combining scRNA-seq with *pax8::GFP* transgenic labeling and targeted *in situ* hybridization, we spatially mapped transcriptionally defined populations within the intact mesonephros. Nephron progenitor cells marked by *six1* localize near the dorsal midline, proximal segments extend laterally and comprise most of the nephron mass, and distal/principal/ionocyte populations converge dorsally into the mesonephric duct. This spatial framework aligns with pseudotime-inferred developmental trajectories and provides anatomical context for lineage branching events.

### Conclusions and evolutionary implications

Together, these findings establish a comprehensive cellular atlas of the *Xenopus* mesonephros and demonstrate that conserved nephron patterning programs operate across successive kidney forms and vertebrate lineages. Our data support a model in which core epithelial segment identities are evolutionarily stable, while intermediate states and lineage branching dynamics are subject to temporal and regulatory modulation. This modular reuse of conserved transcriptional programs may represent a general principle of vertebrate organ evolution, enabling morphological diversification while preserving essential functional architectures.

## Acknowledgements

This work was performed as a joint venture between Dr. Sergei Sokol (Ichan School of Medicine at Mount Sinai), and R. K. Miller. This work was primarily done in the lab of Dr. Sokol under his tutelage, and we thank him for all the work he and his lab invested in this project.

We appreciate the helpful suggestions and advice throughout this project from the members of the laboratories of P. D. McCrea, J. Park, and M. Kloc. Special thanks to Dr. Yoshihiro Komatsu for the use of equipment in this project. We thank the animal care technicians and veterinarians, including J.C. Whitney T.H. Gomez and Keiji Itoh, who took care of the animals. Also, the NXR for providing cryosperm RRID:SCR_013731. Thanks to the UTHealth Cancer Genomics Center (CGC) for the quick processing of the scRNA-seq samples. Additional thanks to the Soeren S Lienkamp, and Oliver Wessely labs for providing the many plasmids for making *in situ* probes. Brandy L. Walker M.S. hand drew artwork for figure 6.

## Competing Interests

‘No competing interests declared’

## Data availability statement

The single-cell RNA sequencing data generated in this study have been deposited in the NCBI BioProject database under accession number PRJNA1274030 (https://dataview.ncbi.nlm.nih.gov/object/PRJNA1274030?reviewer=u32abs94s664qs8ivpt4l6vi1k). Processed expression matrices and metadata are included within the repository. Additional data supporting the findings, including imaging datasets and analysis scripts, are available from the corresponding author upon request.

## Funding

Primary funding for this project was NIH grant 5R35GM122492-08 to Sergei Y. Sokol and private funding from Arkady Volozh. Additional help was provided by National Institute of Diabetes and Digestive and Kidney Diseases, Grant/Award Numbers: R03DK118771 and R01DK115655 to RKM

## Author contributions

RKM and SYS initially conceptualized this project. Kidney dissections and single-cell analysis was performed by MEC. Data analysis was performed by MEC with the help of MA, and NL. Writing and figures were initially assembled by MEC and AR. Project administration as well as data and manuscript evaluation were carried out by MEC, RKM and SYS. *In situ* analysis was carried out by MEC and AR.

## Notes

### Competing Interest Statement

The authors have declared no competing interest.

### Summary of Updates

The author list and contributions have been changed to better represent the work done. Adrian Romero is now co-first Author. Changes to the intro have been made to better reflect the project's goals.

