## Supplementary information for "A single-cell transcriptomic map of the *Xenopus* mesonephros reveals conserved nephron patterning across vertebrate kidney forms"

### Supplemental Figure 1

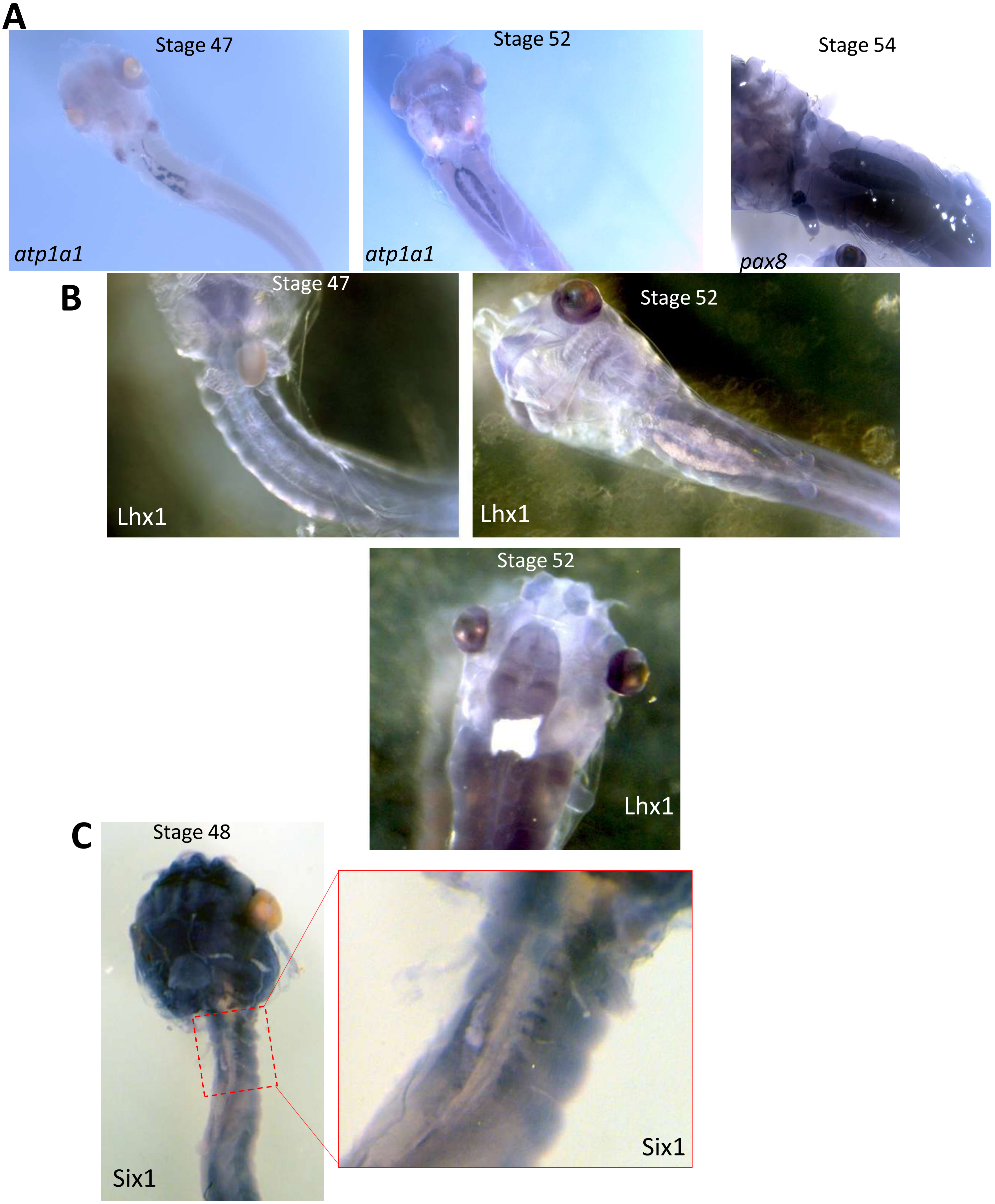


**Figure S1**: A.) Insitu analysis of kidney nephron markers atp1a1, pax8. B.) *lhx1* in situ of stage 47 and 52 kidney and brain staining. C.) Staining of the kidney progenitor marker *six1*.

### Supplemental Figure 2

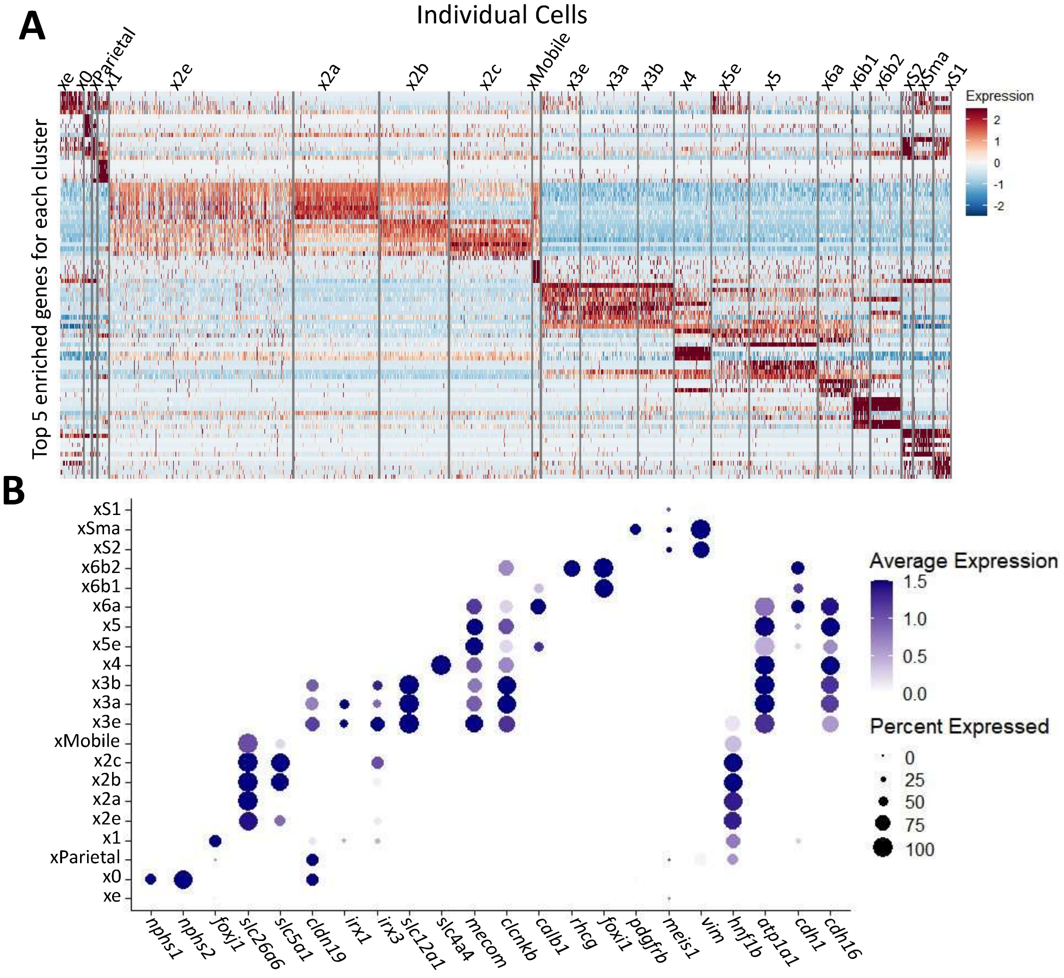


**Figure S2:** A.) Heatmap showing the top 10 genes for each cluster after combining homeologs. Each row indicates expression of an individual gene while each column indicates a single cell. Dotplot showing expression of kidney segment genes.

### Figure Supp 3

#### Non-Kidney Cell Types

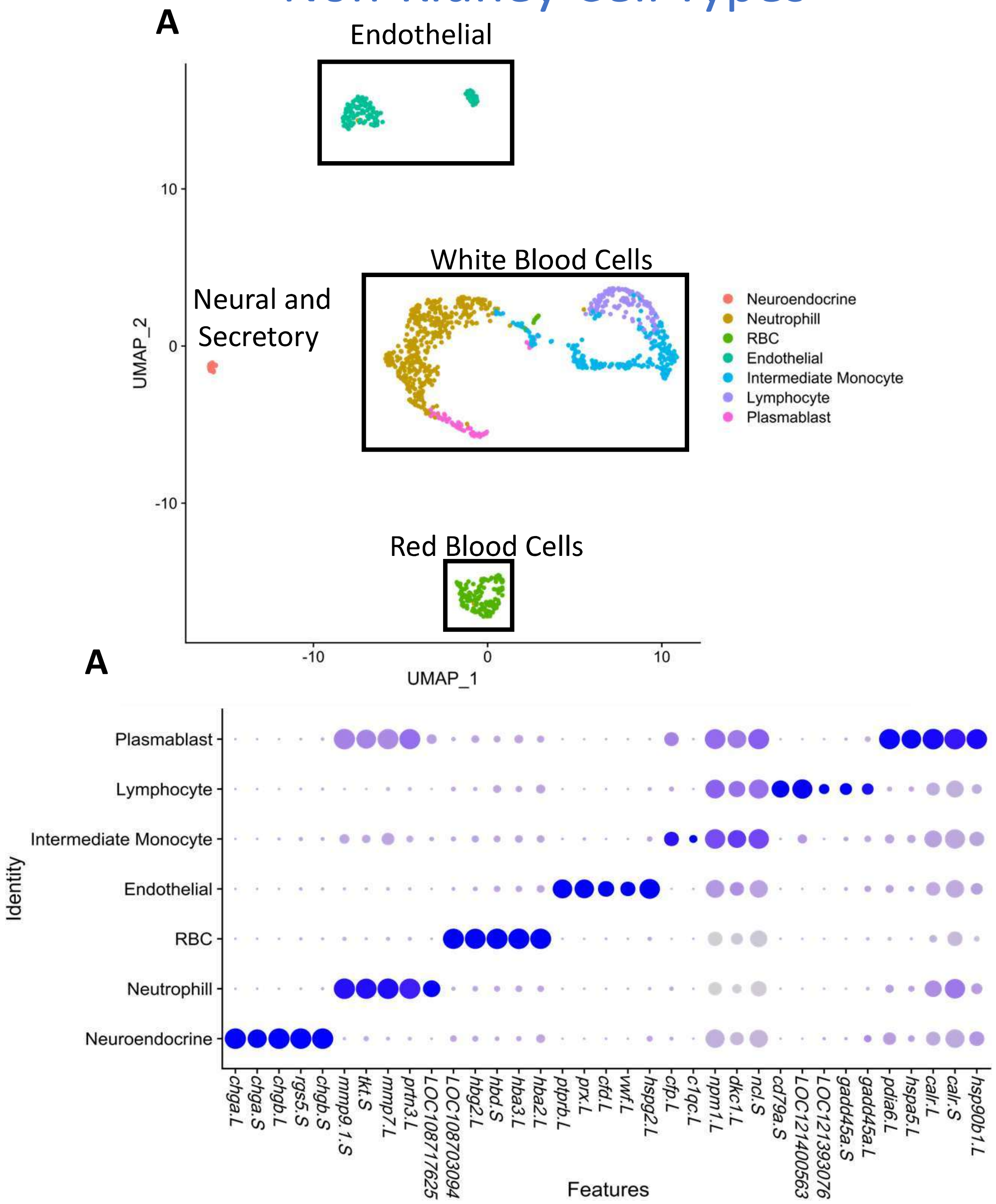


**Figure S4** – Non-kidny cells were isolated and identified. **A)** UMAP projection of non-kidney cells. **B)** Dotplot showing top 5 enriched genes for each cell types.
